# Environmental DNA Metabarcoding Reveals Winners and Losers of Global Change in Coastal Waters

**DOI:** 10.1101/2020.10.08.331694

**Authors:** Ramón Gallego, Emily Jacobs-Palmer, Kelly Cribari, Ryan P. Kelly

## Abstract

Studies of the ecological effects of global change often focus on one or few species at a time. Consequently, we know relatively little about the changes underway at real-world scales of biological communities, which typically have hundreds or thousands of interacting species. Here, we use monthly samples of environmental DNA to survey 222 planktonic taxa along a gradient of temperature, salinity, dissolved oxygen, and carbonate chemistry in nearshore marine habitat. The result is a high-resolution picture of changes in ecological communities using a technique replicable across a wide variety of ecosystems. We estimate community-level differences associated with time, space and environmental variables, and use these results to forecast near-term community changes due to warming and ocean acidification. We find distinct communities in warmer and more acidified conditions, with overall reduced richness in diatom assemblages and increased richness in dinoflagellates. Individual taxa finding greater suitable habitat in near-future waters are more taxonomically varied and include the ubiquitous coccolithophore *Emiliania huxleyi* and the harmful dinoflagellate *Alexandrium sp*. These results suggest foundational changes for nearshore food webs under near-future conditions.

## 4 Main Text

### 4.1 Background

As ocean acidification and warming continue apace, changes in the marine environment are having an effect on many species’ metabolism, development, growth and reproduction success [37, 20, 5, 13], very likely altering food webs [58, 10, 27] and species’ interactions in ways that are poorly understood. Laboratory or mesocosm-based manipulation experiments have documented a wide variety of biological responses under projected climate scenarios of *p*CO_2_, pH, solar radiation, salinity and temperature [23, 16, 40], showing an array of species-specific responses among particular taxa of interest. However, information regarding multi-species or community-wide responses to these stressors is far more limited [38, 32]. The scarcity of such data is likely attributable to the difficulty of simultaneously tracking the responses of many species in the field, and to the difficulty of identifying natural systems that adequately reflect the environmental gradients under study.

Two natural CO_2_ seeps in nearshore marine habitats – one in Italy and one in Papua New Guinea – have demonstrated shifts in benthic communities associated with especially acute acidification in the present day, previewing those we might expect at a more global scale under future conditions [38, 21]. But beyond these exceptional sites, it is difficult to measure changes in ecological communities associated with the relatively subtle shifts in nearshore ocean chemistry observed to date, particularly in light of naturally large spatial and temporal variation in these communities. The Puget Sound in Washington, USA, offers a gradient of carbonate chemistry parameters and other environmental conditions in close geographic proximity. Complex bathymetry, water circulation patterns, and nearshore landforms create intertidal sites exposed to large variations in temperature, *p*CO_2_, pH, and related parameters [36], creating an opportunity to test the effect of these measures on marine communities under conditions expected worldwide in the near future [50], and time-series sampling across the spatial gradient lets us control for site- and season-specific effects. This study system therefore provides a powerful means of modeling community-level responses to changing environmental conditions.

Even given the appropriate environmental gradients, tracking the biological responses of many taxa simultaneously remains challenging. Environmental DNA (eDNA) metabarcoding [29, 33] addresses this problem by amplifying a common gene region out of DNA present in a water sample; the technique can detect hundreds to thousands of taxa per sample, potentially with species-level identification. A growing body of evidence supports the efficacy of eDNA metabarcoding for monitoring biodiversity (see a review in [59]), and this approach has been successfully used to detect community composition variation across environmental changes in aquatic [19], estuarine [12, 41], and marine ecosystems [6, 18].

Here we use series of metabarcoding samples taken across space and time to track changes in nearshore ecological communities associated with differences in pH, water temperature, and other environmental variables. We use broad-spectrum PCR primers [43] to target eukaryotes specifically, identifying the likely effects of future climate scenarios on suites of planktonic taxa.

## 5 Methods

### Sampling

We collected water samples to assess eDNA communities in two regions of the Salish Sea (Washington, USA): San Juan Island and the Hood Canal. These sites experience substantial variation in water chemistry and other environmental conditions despite geographic proximity (ca. 300km; Figure 1). We sampled eight sites monthly for approximately 1.5 years (March 2017 to August 2018), taking three 1L samples (biological replicates; ca. 10m apart) each month at each site (261 bottle samples total). Each sample was filtered through a 0.45 *μ*m cellulose filter, and the filter preserved in Longmire buffer until DNA extraction [56]. Concurrently, we collected one 120 ml water sample from each site and poisoned it with 0.1 ml of saturated HgCl2 for carbonate chemistry analysis, following [57]. We also collected *in situ* measurements of temperature, salinity and dissolved oxygen using a handheld multiprobe (Hanna Instruments, USA) and a portable refractometer. We note that many unmeasured variables influence planktonic communities (e.g., nutrients, sunlight, wave energy), but that our set of measured parameters clearly distinguished communities and was adequate for our purposes.

**Figure 1:**
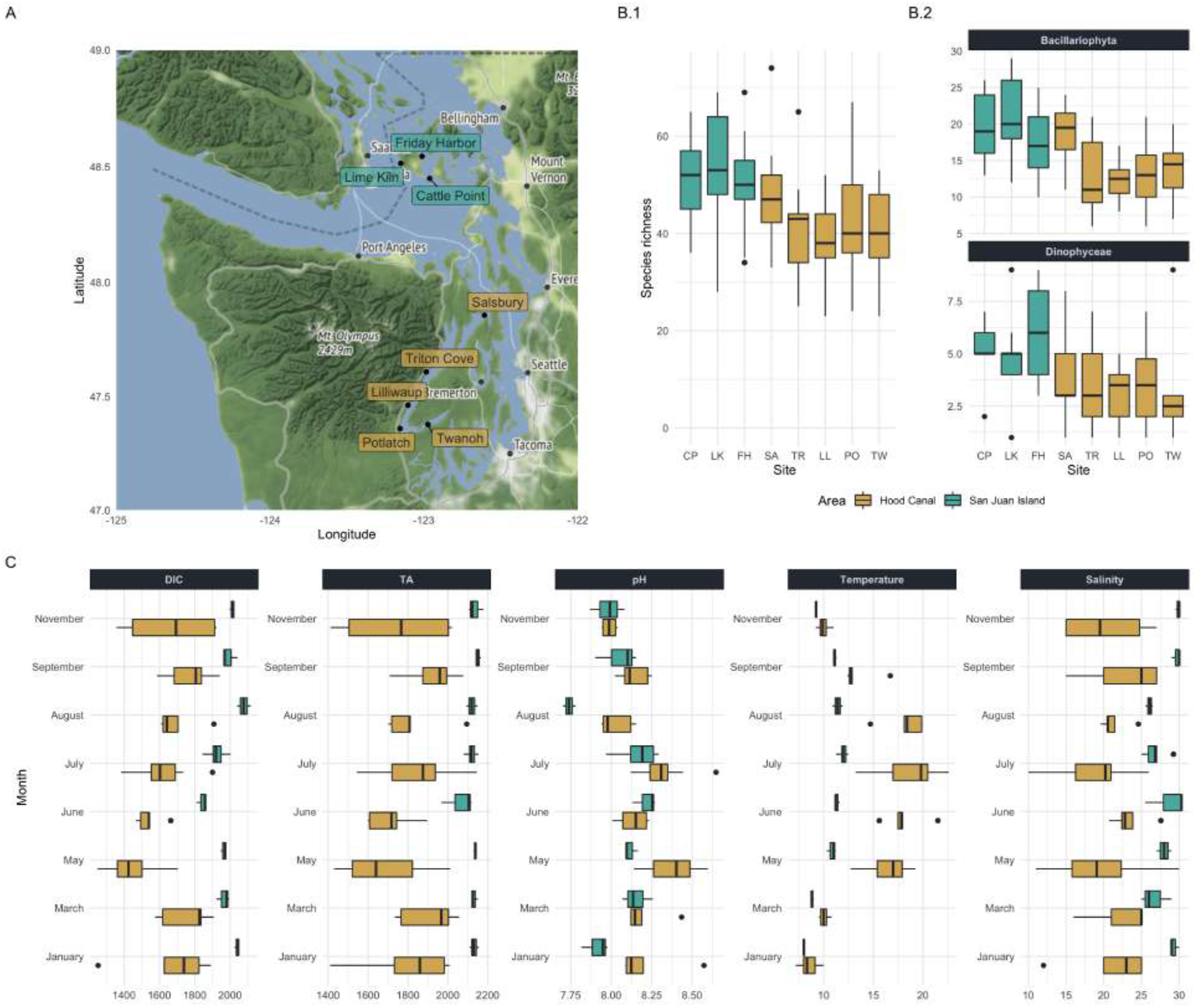
**A** Sampling locations in intertidal areas of the Hood Canal (dark gold) and San Juan Island (turquoise). **B** Planktonic richness (unique taxa) per sampling locality, as reflected by the eDNA COI assay. Boxplots represent the variability in richness across all time points for samples taken at the indicated site. **B.1** Taxa from all phyla; **B.2** Diatoms(above) and dinoflagellates (below) shown separately; note change in the scale of the y-axis. **C** The observed environmental profiles of these two regions reflect a broad range of environmental conditions, with the Hood Canal resembling future conditions in temperate areas worldwide. Shown: Dissolved Inorganic Carbon (DIC, *μ*moles/kg); Total Alkalinity (TA, *μ*moles/kg); Temperature (°C) and Salinity (PSU).

We characterized sample carbonate chemistry by measuring Total Alkalinity (TA; open-cell automated titration based on a 876 Dosimat plus (Metrohm AG) as part of a custom system assembled by Andrew Dickson (UCSD) and used in the laboratory of Alex Gagnon at UW) and Dissolved Inorganic Carbon (DIC; Apollo Instruments, USA; CO_2_ extraction system with 10% (v/v) phosphoric acid). Both measurements were calibrated and validated with certified reference material from the Scripps Institution of Oceanography. Using DIC and TA, we calculated pH and the remaining carbonate system parameters using the R package ‘seacarb’ [25].

Our sampled areas differed in the environmental variables driving changes in carbonate chemistry. San Juan Island was less seasonally variable than the Hood Canal in every measured parameter (Figure 1C); the island is more directly affected by summer coastal upwelling as a function of bathymetry and circulation patterns [50], and this appears to be the dominant influence on carbonate chemistry there. By contrast, photosynthesis and respiration likely drive much of the carbonate chemistry variation in the Hood Canal (See Supplementary Information).

### eDNA sequencing and bioinformatic processes

We purified DNA from each filtered sample using a Phenol-Chloroform-Isoamyl Alcohol protocol, following [56]. After reducing inhibition via a 1/10 to 1/100 dilution, the extract was used as template for a PCR reaction targeting a 313bp fragment of cytochrome oxidase I [43]. PCR reactions were performed in triplicate and sequenced individually to quantify the stochasticity of PCR reactions on a mixed template sample, and we attached secondary indexing tags using a two-step PCR process [51]. PCR conditions and protocols for sample identification followed [34], and batches of 49 to 178 multiplexed samples were sequenced using MiSeq v2-500 or v3-600 sequencing kits using manufacturer protocols. On each sequencing run, we added triplicate samples consisting on DNA obtained from species not present in the marine environment under study (Red Kangaroo (*Macropus rufus*) and Ostrich (*Struthio camelus*)) to establish quality controls of sample assignment and to quantify levels of ‘tag-jumping’ or sample-cross-talk [60].

Code for all quality-screening and bioinformatics is available in the Supplementary Information, implemented in Unix and R [55]. Briefly, we used a Unix script that calls secondary programs for primer-trimming and preliminary quality-control [46, 11] we estimated the likely composition of each sample using DADA2. This approach avoids clustering, such that we retained all of the amplicon sequence variants (ASVs, *i.e*., unique sequences); we subsequently carried out secondary quality-control and decontamination following [34]. We then assigned sequences to known taxa using phylogenetic tree placement with *insect* v1.1 [67]; where *insect* could not place individual taxa, we supplemented assignment by classification against a custom COI database using *anacapa* [15] and *bowtie2* [42]. We conservatively kept only taxa annotated at the level of taxonomic family, genus, or species, so we could reliably infer taxon natural history under the assuming that taxa within the same family shared broad natural-history characteristics. Using published literature and online databases, we placed every recovered taxon into a benthic/planktonic category and focused our analysis on the planktonic community (see Supplementary Information).

By treating amplification efficiency as consistent within a given taxon, we created an index of abundance for each taxon across space and time (“eDNA index” [35]), using pooled data from technical replicates and mean proportions across biological replicates. We used this index of abundance in the multivariate community analysis, and used binary (presence/absence) data to capture species-level responses to environmental conditions. In the Supplementary Information, we provide a function in R to calculate this index.

### Present scenario community analyses

We measured community changes across environmental space using multivariate analyses. We used the index of eDNA abundance to calculate Bray-Curtis dissimilarities between samples, and estimated the effects of temperature, pH, and salinity on community composition using Constrained Analysis of Principal Components (CAP, [3]; ‘capscale’ function in the R package *vegan* [52]).

Independent of environmental parameters, we separately clustered samples by pairwise Bray-Curtis dissimilarities (k-means; N = 3) to identify groups of samples that were similar to one another with respect to biological community. The SIMPER procedure in *vegan* revealed the taxa most strongly contributing to between-cluster differences.

For community-level projections, we coded community-cluster identity (Figure 2) as an unordered response variable in a multinomial logistic model, with temperature, pH, and area (Hood Canal vs. San Juan Island), as predictor variables. Salinity is predicted to remain largely unchanged in future scenarios [36], and because salinity was correlated with temperature in our dataset, it was not an important predictor variable and we subsequently dropped it from our models. We calculated the probability of each community, given these predictors, using a multinomial logistic regression in the R package *nnet* [63].

**Figure 2:**
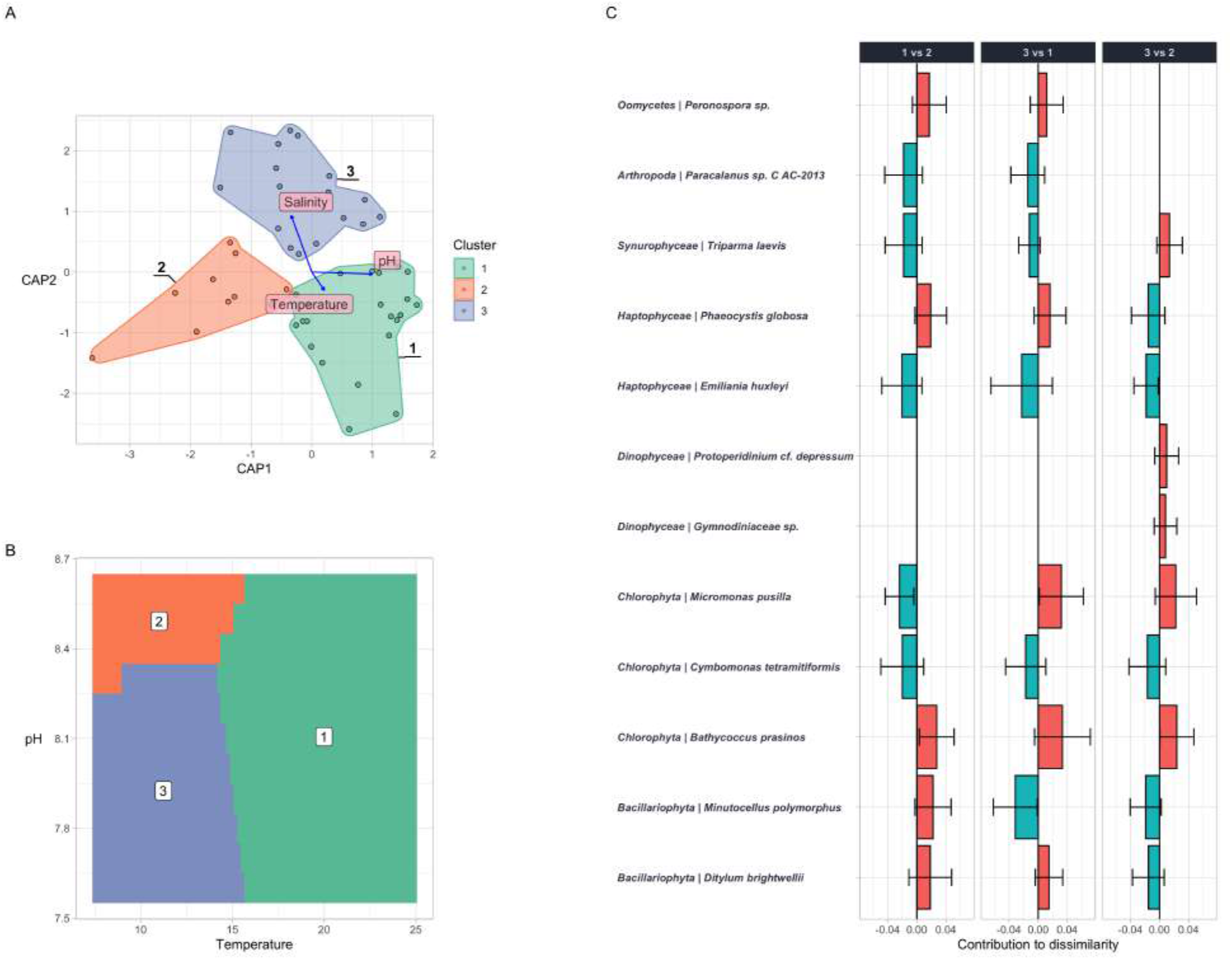
Biological communities and their relationship with environmental variables in the summer months of the Hood Canal. **A** Constrained Analysis of Principal Coordinates (CAP) of Bray-Curtis dissimilarities among biological communities, as constrained by pH, temperature and salinity (arrows). **B** Most-likely cluster as a function of temperature and pH given a multinomial logistic model (salinity was uninformative, being tightly correlated with temperature). **C** Relative abundance (eDNA index; see [35]) of the taxa best distinguishing the three communities illustrated (SIMPER analysis). See Supplementary Information for full analysis.

### Year-2095 Environmental Scenario and Biological Responses

We estimated the distribution of environmental parameters for the overall Salish Sea in 2095 from the results of [36], which estimated an annual mean increase in temperature of 1.51 °C and mean pH decrease of 0.18 for the Salish Sea as a whole. We fit a normal distribution to our 2017 environmental observations to create baseline conditions, and modeled the change in mean parameter values between 2017 and 2095 as a linear function of time. We then used the modeled distribution of environmental parameters to generate 1000 simulations of each year scenario. The scenario labeled as 2095 is the set of parameters falling within the 95% percentile in simulations for the years 2091-2095. See the R code in the Supplementary Information (lines 98-135).

To model biological responses to present and future scenarios, we used a hierarchical logistical regression model relating the presence of each taxon to temperature and pH, in which the slopes of temperature and pH effects varied by taxon, and each taxon had a unique intercept that was allowed to vary by geographic area. For each taxon, we fit these models using the Bayesian generalized linear mixed effects functions in R package rstanarm [26] for R. Model selection using WAIC [65] supported this as the preferred model over several similar ones (see Supplementary Information for model comparison information and code) and helped to avoid model overfitting and maintain out-of-sample predictive power.

Given the sea-surface temperatures and pH values for 2017 (observed) and 2095 (estimated) and taxon-specific logistic regression models, we then evaluated the suitability of habitat for each taxon in the future scenario. For each point in the pH – temperature grid, we calculated a species’ probability of presence as the mean of 100 independent draws of the posterior model response. For each point, the sum of mean probabilities across species provided richness estimations. The mean value across the 100 draws was the input for a Wilcoxon test of differences in species richness between the 2017 and 2095 scenarios in each region. We performed the Wilcoxon test globally (total species richness) and on each phylum independently.

We can only model responses of taxa present in our data set. That is, we may predict that the number of (for example) diatom species present will decline relative to those present today, but our data do not allow us to predict whether new species will immigrate from elsewhere or how species might evolutionarily adapt to future conditions. It is beyond the scope of our work to account for the latter, and furthermore, because of the extreme uncertainty of evolutionary responses, the predictions of species distribution models are often interpreted without considering adaptation or phenotypic plasticity [48].

All raw data used in this manuscript and the scripts that generate the final figures and analysis can be found at https://github.com/ramongallego/eDNA.and.Ocean.Acidification.Gallego.et.al.2020

## Results

### Variation in Carbonate Chemistry and in Ecological Communities

Despite geographic proximity and similar overall species composition (127 of the 222 planktonic taxa were found in both regions and accounted for 98% of the sequences), the areas under study – San Juan Island and the Hood Canal – varied widely in pH, temperature, and other environmental parameters (Figure 1C), with a smooth gradient in conditions along the Hood Canal, and San Juan Island more closely resembling full marine conditions. Different points along the environmental continuum simultaneously showed differences equivalent to those predicted between present-day and future oceans [9].

Metabarcoding analysis of eDNA samples generated more than 50.8M sequences from 778 samples. These samples represented biological and technical replicates from 86 unique sampling events. After bioinformatics quality-control the dataset included ~45M sequences, from 4849 unique amplicon sequence variants (ASVs). Of these, 1364 ASVs (22.6M reads) could be annotated to a taxonomic level of Family or lower. These ~ 500 taxa from 43 phyla were split according to their natural history and habitat (benthic vs. planktonic; see Supplementary Information). Because we expect planktonic taxa to vary with water mass [34] and therefore with bottle-sampled carbonate chemistry, here we focus on only the planktonic taxa (N = 221). These taxa showed a seasonal richness gradient between study areas, consistent with documented biodiversity clines in the area (Figure 1B.1) [17].

Bray-Curtis dissimilarities among samples revealed large differences in metabarcoding communities due to geographic Area (Hood Canal vs. San Juan Island; F = 1.6184; p < 0.01). We therefore performed a constrained analysis of principal components (CAP) for samples within each Area, showing the differences among communities as a function of temperature, pH, and salinity (Figure 2; for clarity, results for Hood Canal shown; full analysis in the Supplementary Information).

Each biological cluster (colored hulls, Figure 2A) occupied a unique area of environmental parameter space. Planktonic communities therefore varied predictably with water temperature, salinity, and pH, across a range of those parameters likely to be encountered in many near-term future-ocean scenarios [50]. Multinomial logistic regression yielded predictions of the most-likely community for any combination of environmental parameters (Figure 2B).

These clusters were distinguished by changes in the relative abundances of a wide variety of taxa. In the Hood Canal for example, the cluster linked with colder water and higher-pH (cluster 2 in Figure 2B) showed higher eDNA indexes of diatoms (*Minutocellus polymorphus*), green algae (*Bathycoccus prasinos* and *Micromonas pusilla*), and dinoflagellates like *Karlodinium* sp. relative to the lower-pH cluster at the same temperature range. In that community (cluster 3) *Emiliania huxleyi, Ditylum brightwellii* and the copepod *Paracalanus* sp. C are more prevalent. The cluster common in the high temperature range (cluster 1 in Figure 2B) shows high values of *Phaeocystis globosa,Emiliania huxleyi, Ditylum brightwellii* among other species. Cluster 1 (Figure 2A,B) and similar planktonic communities occupy the spectrum of environmental conditions most likely to be encountered in near-future climate scenarios as temperature rises and pH falls. For example, we expect the Hood Canal in 2095 to have the conditions in which community 1 is the most likely community 67% of the time, an increase of 11% compared to 2017(see Supplementary Information). On the other hand, conditions that make cluster 2 the most likely community drop from 12% to 2% of the time between 2017 and 2095.

### Climate envelopes and future distributions

To explore the suitability of different environmental conditions for each taxon, we modeled the likelihood of taxon presence as a function of temperature, pH, and geographic Area as described in the Methods. Salinity was not informative in our models, as it is highly correlated with temperature in our dataset and is moreover predicted to remain largely unchanged in future scenarios [36]. Model projections let us show the change in the probability of presence of each individual taxon for 2095 vs 2017 (Figure 3 A.1, B.1, C.1), estimate richness for larger taxonomic grounds as a whole for these two climates (Figure 3 A.2, B.2, C.2), and estimate richness within taxonomic groups across the pH-temperature continuum (Figure 3 A.3, B.3, C.3).

**Figure 3:**
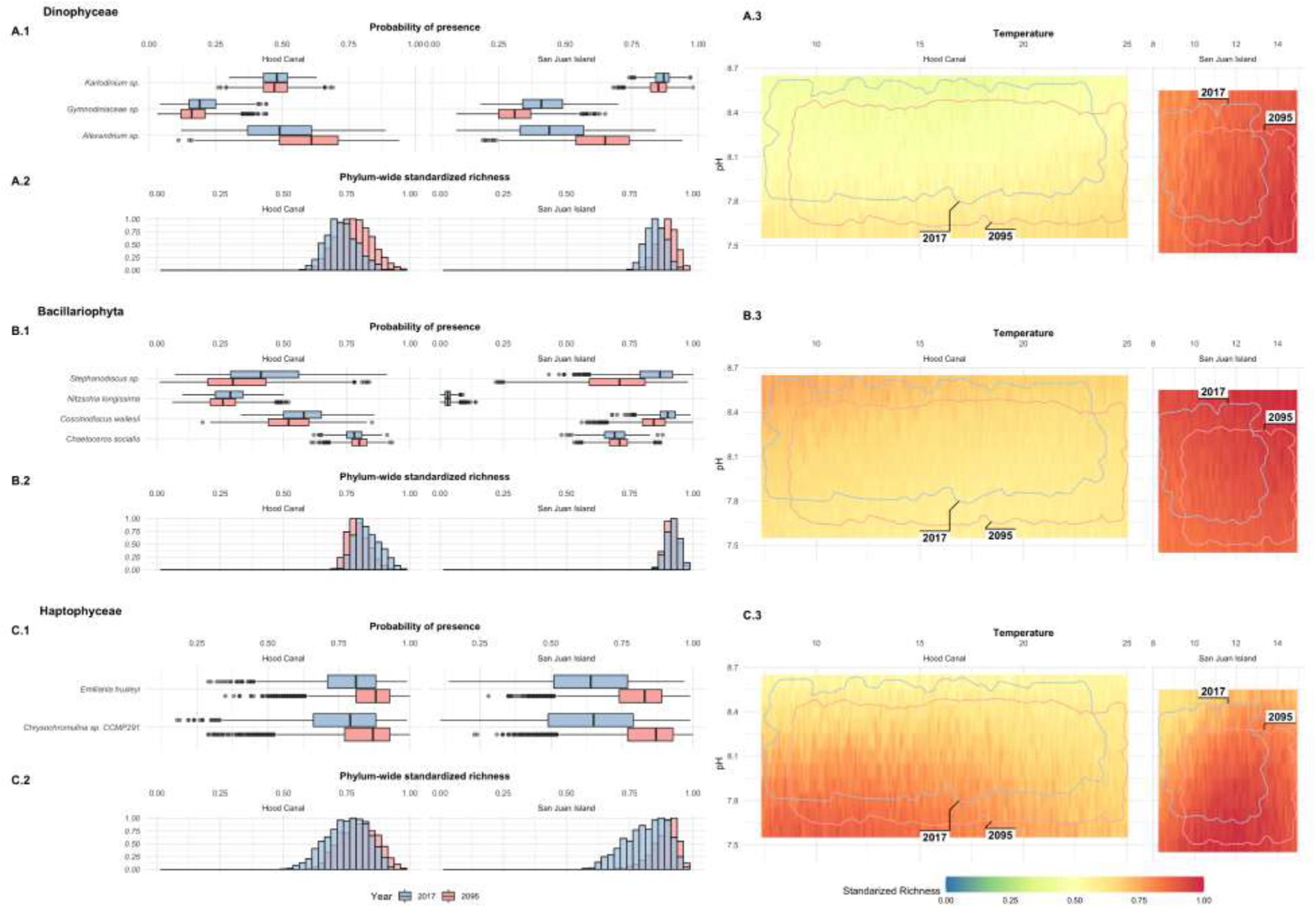
Forecasted changes in plankton in the Salish Sea for Dinoflagellates (panel **A**), Diatoms (**B**) and Haptophytes (**C**). Each panel shows: (**1**) probability densities for the occurrence of selected taxa (species- and genus-level) for 2017 (blue) and 2095 (red); data are mean probabilities over 100 model draws, and variance in probability is due to differences in underlying environmental conditions. (**2**) Changes in relative species richness within each phylum across the simulated scenario. (**3**) Relative taxon richness (raster color, warmer colors are more taxon-rich) for each of these same higher-taxa, for plausible ranges of pH and Temperature. Envelopes of observed (2017, blue) and modeled (2095, red) annual conditions in the Salish Sea shown for reference. Hood Canal and San Juan Island plotted separately to illustrate environmental differences between them.

Diatoms (Bacillariophyta) show the steepest richness decline under future conditions (Figure 3 B.2); the probability of occurrence decreases markedly for diatom taxa including *Coscinodiscus* and *Stephanodiscus*, both centric body forms. These declines in diatom richness were more accentuated at lower pH values and higher temperatures. Other taxa likely to find less-suitable habitat in the future include the dinoflagellate Gymnodiniaceae sp.

Likely winners under future conditions are more widely scattered among higher taxonomic groups. The haptophytes *Emiliania huxleyi* and *Chrysochromulina* sp. the dinoflagellate *Alexandrium* sp. all find more suitable habitat in both of our study areas. Among others not shown in Figure 3, *Chaetoceros* (diatom) and many hydrozoans (Cnidaria) likely increase in San Juan Island, and the potentially fish-killing heterokont flagellate *Pseudochattonella* increases in both study areas. See Supplementary Information for a complete list of taxon-specific projections.

Given such heterogeneity in projections, gains and losses tend to balance one another out when looking at overall richness variation; we find no change in median richness for the year 2095 relative to the present in the Hood Canal (overall taxon richness by year, 95% confidence interval in median species richness −0.08, +0.04; Wilcoxon p = 0.5); while higher diversity is expected in the San Juan Island in 2095 (increase in median species richness of 1.84-2.2, p < 10^−16^). Diatoms, in particular, show small but significant declines in richness in the Hood Canal (0.46-0.55 species, p < 10^−16^), while the changes on the San Juan Island are negligible (median change 0-0.08 p = 0.04). Dinoflagellates see their richness increase in both regions with the future scenario (median change 0.14-0.18 Hood Canal; 0.21-0.25 San Juan Island; p < 10^−16^ for each).

The bulk of our projected community changes result from now-rare conditions occurring more frequently in the future. For example, in in the Hood Canal at present, we expect surface waters to have pH < 7.9 and T > 19°C only 1% of the time. In 2095, we expect these conditions 6 times more frequently (i.e., 6% of the time). At these values of T and pH, our model predicts the harmful *Alexandrium* sp. to occur more often than not (mean frequency of occurrence = 0.83). By contrast, the large centric diatom *Coscinodiscus* – a potentially key source of carbon for zooplankton and small fishes [53, 69] with effects on dissolved oxygen and other water-column characteristics [44] – occurs only one-third of the time under these same conditions (mean frequency = 0.35).

## Discussion

Temperate surface oceans worldwide average approximately 14°C and pH of 8.1 [9], and will change substantially in this century [mean ΔT 2.5°C, ΔpH −0.35 globally; RCP 8.5; 24]. Here we document communities exposed to this same range of projected conditions in the present day, along an environmental gradient only ca. 200km wide, allowing us to project future ocean communities from a robust set of underlying observations. Our results reflect patterns in a diverse selection of species from nearshore marine communities in the Salish Sea, consisting of 222 planktonic taxa obtained from the metabarcoding analysis of 227 discrete samples across 77 space-time points (eight sites, 1.5 years). We find that changes in the composition of biological communities closely mirrored the variation in pH and temperature, with clear winners (e.g., *Emiliania huxleyi, Alexandrium*, and others) and losers (many, but not all, diatoms) likely to shift the structure and function of future marine communities.

A vast amount of evidence suggests climate-associated effects on marine species, and broad patterns of sensitivity are discernible within major taxonomic groups [24, 61, among many others]. However, because the strength and direction of these effects are variable and species-specific [39], very little is known about community-level impacts. Our work illustrates the nearshore planktonic communities that can thrive in low pH – high temperature conditions; such communities are therefore likely to become more prevalent under future conditions.

The large number of species and broad set of environmental conditions we sampled yield substantial inferential power despite lacking the the degree of experimental control present in a laboratory or mesocosm.

Among the taxa surveyed, diatoms are of particular interest for their ubiquity in the world’s oceans and their important roles in marine food webs [4, 64], as well as in ecological and evolutionary theory [45]. Our model suggests that diatoms will decrease in richness between the present and 2095, particularly in the Hood Canal, where extreme temperatures are more common. Although the most prevalent response among diatoms is a decrease in suitability, some substantial variability in responses exists within the group. For example, the centric diatom *Coscinodiscus* spp., which is a food source for *Acartia* spp. copepods [31] and many other animal species, will see future suitable habitat only in colder waters such as those in San Juan Island, while *Skeletonema* spp. and the harmful algal bloom (HAB)-forming species *Pseudo-nitzschia* spp. will see their habitat suitability remain constant or slightly increased, especially at low pH levels (see Supplementary Information).

More strikingly, we see a dramatic increase in suitable environment for the HAB-forming dinoflagellate *Alexandrium* sp., which can substantially harm local ecosystems [14] and economies [1]. This increase is particularly high in the summer months of the Hood Canal, when pH is low and temperatures are are high. Both archaeological and experimental evidence suggest *Alexandrium* sp. blooms with warmer temperatures [49], and models [48] also predict an increase in bloom-favorable conditions for *Alexandrium* sp. in future oceans.

Our results therefore suggest a possible change in relative dominance between diatoms and other phytoplankton species such as dinoflagellates, consistent with those seen at ecological regime shifts found elsewhere [64, 28]. Such a shift could affect ecosystems in many ways; even under the assumption that the surviving taxa would maintain the primary production levels, for example, the smaller cell-size of dinoflagellates and the differential sinking rates of the two groups would likely alter regional patterns of nutrient cycling and carbon sequestration [68, 2, 7, 54]. Although the north Atlantic has shown an increase in diatom abundance [28], the increase in wind stress and associated mixing in the water column in the open ocean is unlikely to occur in the Hood Canal, where stratification is the strongest in the Salish Sea [47]. Furthermore, locally focused models support an increase in dinoflagellate dominance with climate change, particularly during summer months [36].

Our model also suggests increased environmental suitability for the coccolithophore *Emiliania huxleyi*. There is evidence supporting increased calcification and respiration rates with higher pCO_2_ levels [30] for this ubiquitous species, although the many strains of this species and its adaptive capacity make it difficult to predict longer-term effects with confidence [8].

Changes in environmental conditions and associated shifts in planktonic communities will likely reshape ecosystems and food webs, although some environmental processes may be conserved even as the particular taxa change. A switch from a diatom-dominated ecosystem to one in which dinoflagellate blooms extend in space and time could provoke cascade effects [68] including fish mortality, anoxia [2], and carbon sinking dynamics [7]. Beyond the phylum-specific patterns, the increase in suitable habitat for harmful algae species will alone be an engine for ecosystem change [62, 66].

One general challenge for model-based work is a tendency to extrapolate from observed conditions in ways that are often untestable – by necessity, projections frequently operate outside the range of parameters on which the model was trained [22]. Our study system lets us avoid this pitfall, in that our observed conditions encompass much of the environmental range predicted for future temperate oceans. That is, the changes we predict for the year 2095 do not primarily come from extreme values of pH and temperature, but rather from presently-rare conditions becoming more common.

The taxa surveyed here are a function of our metabarcoding PCR primers [43] and reflect the current status of genetic databases, rather than a complete sampling of the planktonic community; we view these results as a cross-section of common taxa useful for understanding the biological effects of ocean conditions. Our observations are strong evidence of the kinds of changes likely in future marine communities, and they offer testable predictions about the magnitude and direction of effects on focal species.

## Author contributions

RG led sampling, laboratory work, data analysis, and writing. RPK conceived of the project and assisted in sampling, analysis, and writing, and received funding for the work. EJP and KC assisted in sampling, laboratory work, and analysis.

## 6 Acknowledgments

We are grateful to the UW Center for Environmental Genomics and the Conservation Biology Molecular Genetics Laboratory at NOAA Fisheries (Montlake) for access to lab equipment and associated expertise and support. Thanks to Micah Horwith and the Washington State Department of Natural Resources for in-field support, to Terrie Klinger for feedback on an earlier draft, to Stephanie Moore for advice on phytoplankton generally, to Ana Ramoón-Laca for laboratory help, to Dan Drinan for field help, and to Alex Gagnon for access to carbonate-chemistry expertise and equipment. The comments of two reviewers helped us improve the manuscripts and encourage us to streamline the analysis of our data.

